# simuG: a general-purpose genome simulator

**DOI:** 10.1101/491498

**Authors:** Jia-Xing Yue, Gianni Liti

## Abstract

**Summary:** Simulated genomes with pre-defined and random genomic variants can be very useful for benchmarking genomic and bioinformatics analyses. Here we introduce simuG, a lightweight tool for simulating the full-spectrum of genomic variants (SNPs, INDELs, CNVs, inversions and translocations) for any organisms (including human). The simplicity and versatility of simuG makes it a unique general purpose genome simulator for a wide-range of simulation-based applications.

**Availability and implementation:** Code in Perl along with user manual and testing data is available at https://github.com/yjx1217/simuG. This software is free for use under the MIT license.

## 1 Introduction

Along with the rapid progress of genome sequencing technologies, many bioinformatics tools have been developed for characterizing genomic variants based on genome sequencing data. While there is an increasing availability of experimentally validated gold-standard genome sequencing data set from real biological samples, *in silico* simulation remains a powerful approach for gauging and comparing the performance of bioinformatics tools. Correspondingly, many read simulators have been developed for different sequencing technologies, such as ART (Huang *et al*., 2012) for Illumina and 454, SimLoRD (Stöcker *et al*., 2016) for PacBio, and DeepSimulator (Li *et al*., 2018) for Oxford Nanopore. However, when it comes to tools for simulating genome sequences with embedded variants, the choices appear much more limited. The current available tools are either too simple or too specialized. For example, SInC (Pattnaik et al., 2014) can introduce random single nucleotide polymorphisms (SNPs), Insertion/Deletions (INDELs), and copy number variants (CNVs) into a user-provided reference genome but lacks the ability to simulate known variants, which is actually highly relevant in some simulation applications. Simulome (Price *et al*., 2017) is another random variant simulator that provides finer control options, but it is designed for prokaryote genomes only. More sophisticated tools exist, such as VarSim (Mu *et al*., 2015) and Xome-Blender (Semeraro *et al*., 2018), but these tools are mostly tailored for human cancer genome simulation and often require additional third-party databases. Therefore, we feel there is need for a genome simulator that strikes a balance between simplicity and versatility. With this in mind, we developed a general-purpose genome simulator simuG, which is versatile enough to simulate both small (i.e. SNPs and INDELs) and large (i.e. CNVs, inversions, and translocations) genomic variants while staying lightweight with no extra dependency and minimal input requirements. In addition, simuG provides a rich array of fine-grained controls, such as simulating SNPs in different coding partitions (e.g. coding sites, noncoding sites, 4-fold degenerate sites, or 2-fold degenerate sites); simulating CNVs with different formation mechanisms (e.g. segmental deletions, dispersed duplications, and tandem duplications); and simulating inversions and translocations with specific types of breakpoints. These features together make simuG highly amenable to a wide range of application scenarios.

## 2 Description and feature highlights

simuG is a command-line tool written in Perl and supports all mainstream operating systems. It takes the user-supplied reference genome (in FASTA format) as the working template to introduce non-overlapping genomic variants of all major types (i.e. SNPs, INDELs, CNVs, inversions, and translocations). SNP and INDELs can be introduced simultaneously, whereas CNVs (implemented as segmental duplications and deletions), inversions, and translocations can be introduced with separated runs. For each variant type, simuG can simulate pre-defined or random variants depending on specified options. For pre-defined variants, a user-supplied VCF file that specifies all desired variants is needed, based on which simuG will operate on the input reference genome to introduce the corresponding variants. For random variants, simuG supports a wide-spectrum of fine control options, such as ‘-titv_ratio’forspecifyingthetransition/transversionratioofSNPs,‘-indel_size_powerlaw_alpha’ and ‘-indel_size_powerlaw_constant’ for specifying the size distribution of INDELs, ‘-cnv_gain_loss_ratio’ for specifying the ratio of segmental duplication versus segmental deletion, “-duplication_tandem_dispersed_ratio” for specifying the ratio of tandem versus dispersed duplications, and ‘-centromere_gff’ for specifying the location of centromeres so that simulated random CNVs, inversions, and translocations will not disrupt the specified centromeres. An ancillary script vcf2model.pl is further provided to directly calculate the best parameter combinations for the random SNP/INDEL simulation based on real data. Moreover, given the strong association between gross chromosomal rearrangement breakpoints and repetitive sequences (e.g. transposable elements) observed in empirical studies (Zhang *et al*., 2011; Yue *et al*., 2017), simuG can restrict random inversions and translocations to only use user-defined breakpoints (by specifying the ‘-inversion_breakpoint_gff’ or ‘-translocation_breakpoint_gff’ option). The specific feature type and strand information of these user-defined breakpoints will be considered during the breakpoint sampling. For example, the breakpoint pairs that can trigger inversion should belong to the same feature type but from opposite strands (e.g. inverted repeats). Also, when specified, centromeres will be given special consideration in random translocation simulation so that translocations leading to dicentric chromosomes will not be sampled. Finally, when needed, users can also define a list of chromosomes (e.g. mtDNA) to be excluded from variant introduction. Upon the completion of the simulation, three files will be produced: 1) a simulated genome bearing introduced variants in FASTA format, 2) a tabular file showing the genomic locations of all introduced variants relative to both the reference genome and the simulated genome, 3) a VCF file showing the genomic locations of all introduced variants relative to the reference genome. Since simuG’s major input/output formats (e.g. FASTA, VCF, and GFF3) are all widely used in the field, it should be fairly straightforward to connect simuG with other computational tools both upstream and downstream. Please note that when comparing the VCF outputs from simuG and other tools, all VCF files used for the such comparison should be normalized by tools like vt (Tan *et al*., 2015) beforehand.

## 3 Application demonstration

To demonstrate the application of simuG in a real case scenario, we ran simuG with the budding yeast *Saccharomyces cerevisiae* (version R64-2-1) and human (version GRCh38) reference genomes to generate nine simulated genomes for each organism: A) with 10000 SNPs, B) with 1000 random INDELs, C) with 10 random CNV due to segmental deletions, D) with 10 random CNV due to dispersed duplications, E) with 10 random CNV due to tandem duplications, F) with 5 random inversions, G) with 5 random inversions triggered by breakpoints sampled from pre-specified transposable elements (TEs), H) with 5 random translocation, I) with 5 random translocation triggered by breakpoints sampled from pre-specified TEs. Based on each simulated genome, 50X 150-bp Illumina paired-end reads and 25X PacBio reads were simulated with ART (Huang *et al*., 2012) and SimLoRd (Stöcker *et al*., 2016) respectively and subsequently mapped to the yeast and human reference genomes. The read mapping was performed by BWA (Li and Durbin, 2009) for Illumina reads and by minimap2 (Li, 2018) for PacBio reads. With this setup, we evaluated the performance of different variant callers for both small and large variants (Table 1 and Supplementary Note). For small-variants (i.e. SNP and INDELs), we found freebayes (Garrison and Marth, 2012) and the GATK4 HaplotypeCaller (Poplin *et al*., 2018) both performed well, with the latter one marginally won out in INDEL calling. For large structural variants like CNVs, inversions, and translocations, we found both the short-read-based callers Delly (Rausch *et al*., 2012) and Manta (Chen *et al*., 2016) and the long-read-based caller Sniffles (Sedlazeck *et al*., 2018) were able to identify most simulated events, especially when no TEs were associated with the breakpoints. The long-read caller Sniffles showed superior accuracy in resolving the exact breakpoints to the basepair resolution than short-read-based callers by taking advantage of the long-reads, even with half of the sequencing coverage. Between the two short-read-based callers, Manta outperformed Delly in terms of breakpoint accuracy at the basepair level

**Table 1.**
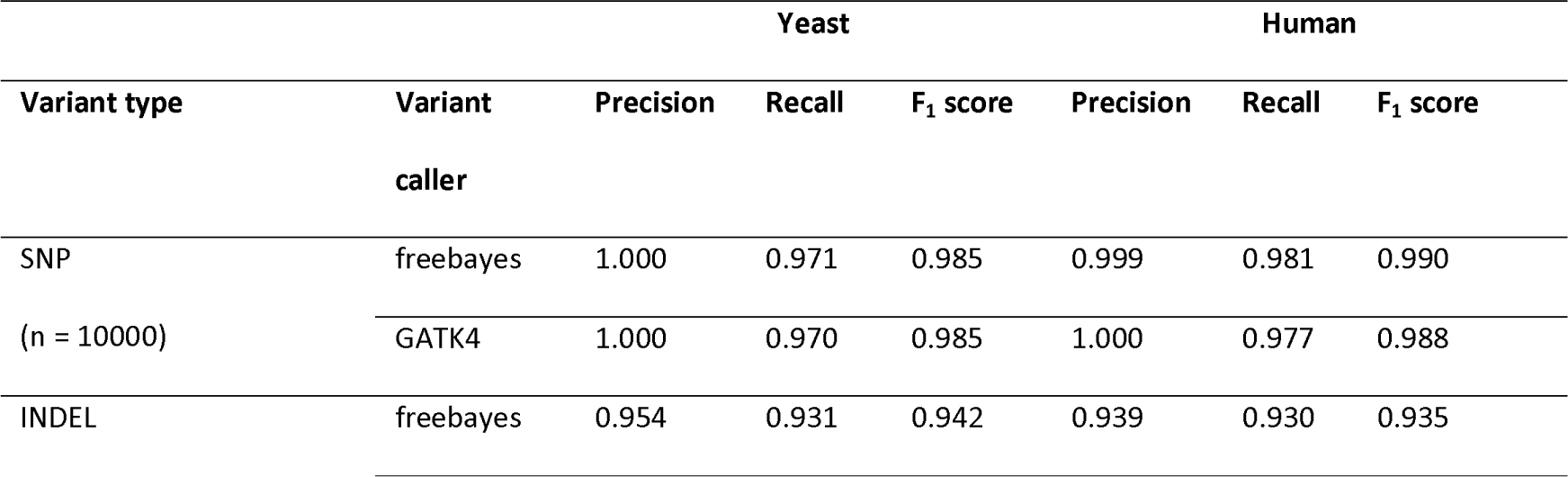
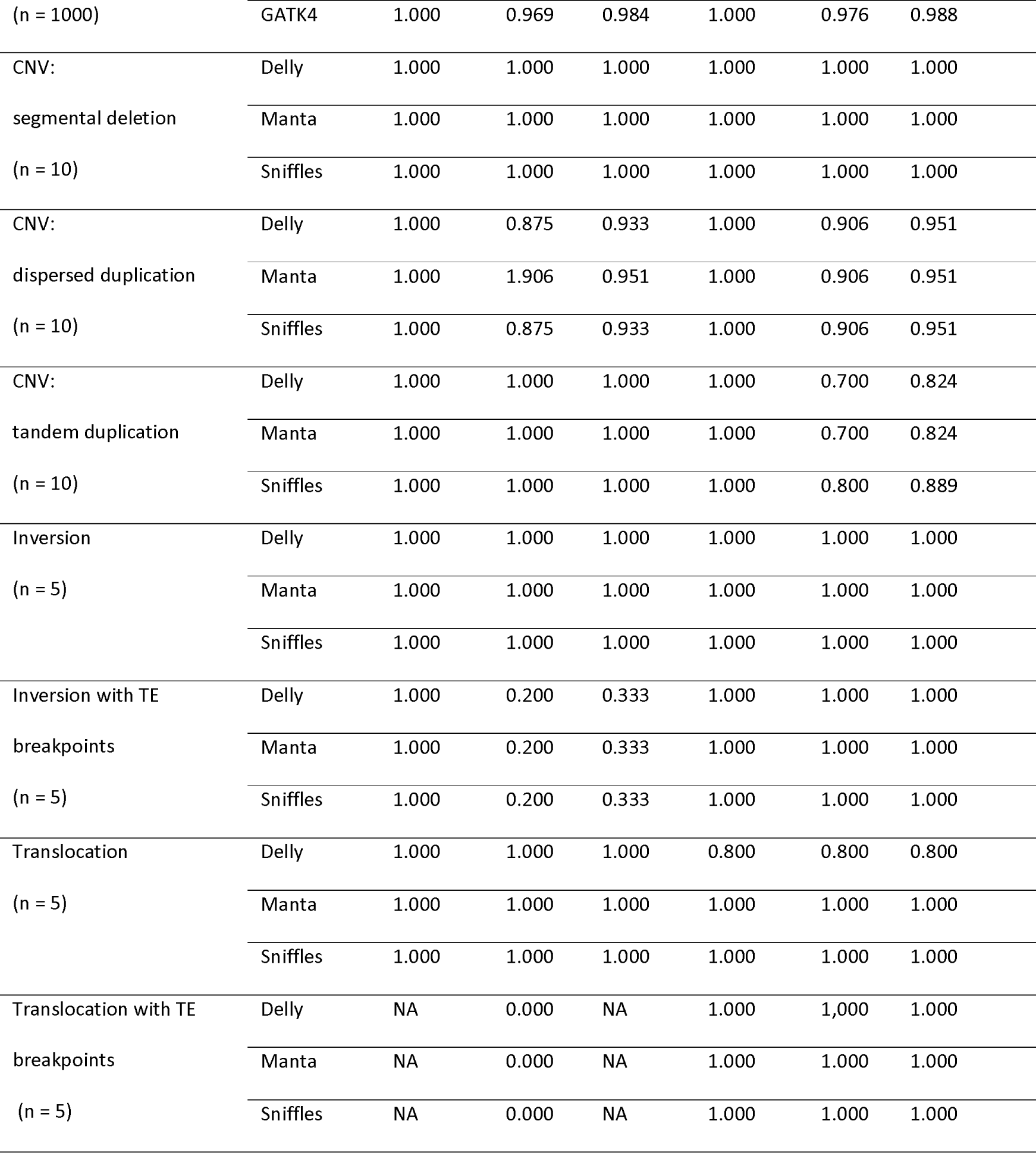
Benchmarking popular variant callers with the small and large genomic variants simulated by simuG. For each variant type, number of introduced variants are shown in parentheses. TE: transposable elements (full-length Ty1 for *S*. *cerevisiae* and full-length intact L1 for human). Precision = true positive/(true positive + false positive). Recall = true positive/(true positive + false negative). F_1_ score = 2 * (recall * precision)/(recall + precision). For a single CNV derived from dispersed duplication, there could be multiple duplicated copies inserted to different genomic locations, making it tricky to calculate accuracy, precision, and F_1_ score by measuring the number of recovered CNV events. Therefore, we calculated these values based on the number of recovered breakpoints instead in this case.

## 4 Conclusions

We developed simuG, a simple, flexible, and powerful tool to simulate genome sequences with both pre-defined and random genomic variants. Simple as it is, simuG is highly versatile to handle the full spectrum of genomic variants, which makes it very useful to serve the purpose of various simulation studies.

