## Supplementary Note for "simuG: a general-purpose genome simulator"

April 29, 2019

**Detailed method description on our variant calling benchmarking analysis**

1. **Input reference genome and annotation**

For our test with the yeast genome, we downloaded the reference genome of the budding yeast *Saccharomyces cerevisiae* S288C (version R64-2-1) as well as the associated genomic feature annotation GFF3 file from the *Saccharomyces* Genome Database (SGD) (https://www.yeastgenome.org/). The centromere and the full-length Ty1 and Ty3 transposable element (TE) annotation were further retrieved based on this GFF3 file.

For our test with the human genome, we downloaded the human reference genome (version GRCh38) from The National Center for Biotechnology Information (NCBI) website (<ftp://ftp.ncbi.nlm.nih.gov/genomes/all/GCA/000/001/405/GCA_000001405.15_GRCh38/seqs_for_alignment_pipelines.ucsc_ids/GCA_000001405.15_GRCh38_no_alt_analysis_set.fna.gz>). The corresponding centromere annotation was retrieved based on the centromere annotation track of the UCSC genome browser (<http://genome.ucsc.edu/cgi-bin/hgTables?hgsid=717837565_0v0BSTaoWRLH5uFpBkbYrt6BnLmm&clade=mammal&org=Human&db=hg38&hgta_group=map&hgta_track=centromeres&hgta_table=0&hgta_regionType=genome&position=chr1%3A1-248%2C956%2C422&hgta_outputType=gff&hgta_outFileName>=). In the case where multiple centromere gaps were annotated for the same chromosome, we treated them as a whole and took their collective outermost boundaries to denote the location of the corresponding centromere. For human TEs annotation, we adopted the full-length intact L1 transposable element annotation from the L1Base2 database (<http://l1base.charite.de/>)

2. **Genome simulation**

With the reference yeast and human genomes as the working templates, we ran simuG with the following parameters to derive each simulated genome:

Simulated yeast genome A (10000 SNPs):

```
perl simuG.pl \
    -refseq SGDref.R64-2-1.fa.gz \
    -snp_count 10000 \
    -titv_ratio 2.0 \
    -excluded_chr_list excluded_chr_list.yeast.txt \ # chrMT excluded
    -prefix yeast_test.SNP_INDEL \
-seed 201812201903
```

Simulated yeast genome B (1000 INDELs):

```
perl simuG.pl \
    -refseq SGDref.R64-2-1.fa.gz \
    -indel_count 1000 \
    -excluded_chr_list excluded_chr_list.yeast.txt \ # chrMT excluded
    -prefix yeast_test.SNP_INDEL \
    -seed 201812201903
```

Simulated yeast genome C (10 random CNVs due to segmental deletions):

```
perl simuG.pl \
    -refseq SGDref.R64-2-1.fa.gz \
-cnv_count 10 \
-cnv_gain_loss_ratio 0 \
    -centromere_gff SGDref.R64-2-1.centromere.gff3 \
    -excluded_chr_list excluded_chr_list.yeast.txt \ # chrMT excluded
    -prefix yeast_test.CNV_DEL \
    -seed 201812201903
```

Simulated yeast genome D (10 random CNVs due to dispersed duplications):

```
perl simuG.pl \
    -refseq SGDref.R64-2-1.fa.gz \
-cnv_count 10 \
-cnv_gain_loss_ratio Inf \
-cnv_max_copy_number 3 \
-duplication_tandem_dispersed_ratio 0 \
    -centromere_gff SGDref.R64-2-1.centromere.gff3 \
    -excluded_chr_list excluded_chr_list.yeast.txt \ # chrMT excluded
    -prefix yeast_test.CNV_DispersedDup \
    -seed 201812201903
```

Simulated yeast genome E (10 random CNVs due to tandem duplications):

```
perl simuG.pl \
    -refseq SGDref.R64-2-1.fa.gz \
-cnv_count 10 \
-cnv_gain_loss_ratio Inf \
-cnv_max_copy_number 3 \
-duplication_tandem_dispersed_ratio Inf \
    -centromere_gff SGDref.R64-2-1.centromere.gff3 \
    -excluded_chr_list excluded_chr_list.yeast.txt \ # chrMT excluded
    -prefix yeast_test.CNV_DispersedDup \
    -seed 201812201903
```

Simulated yeast genome F (5 random inversions):

```
perl simuG.pl \
    -refseq SGDref.R64-2-1.fa.gz \
-inversion_count 5 \
       -inversion_max_size 1000000 \
    -centromere_gff SGDref.R64-2-1.centromere.gff3 \
    -excluded_chr_list excluded_chr_list.yeast.txt \
    -prefix yeast_test.INV_run1 \
    -seed 201812201903
```

Simulated yeast genome G (5 random inversions with breakpoints sampled from full-length Ty1 transposable elements annotated in the reference genome):

```
perl simuG.pl \
    -refseq SGDref.R64-2-1.fa.gz \
-inversion_count 5 \
-inversion_max_size 1000000 \
-centromere_gff SGDref.R64-2-1.centromere.gff3 \
-inversion_breakpoint_gff SGDref.R64-2-1.Ty1_FullLength.gff3 \
    -excluded_chr_list excluded_chr_list.yeast.txt \ # chrMT excluded
    -prefix yeast_test.INV_run2 \
    -seed 201812201903
```

Simulated yeast genome H (5 random translocations):

```
perl simuG.pl \
    -refseq SGDref.R64-2-1.fa.gz \
    -translocation_count 5 \
    -centromere_gff SGDref.R64-2-1.centromere.gff3 \
    -excluded_chr_list excluded_chr_list.yeast.txt \ # chrMT excluded
    -prefix yeast_test.TRA_run1 \
    -seed 201812201903
```

Simulated yeast genome I (5 random translocations with breakpoints sampled from full-length Ty1 transposable elements annotated in the reference genome):

```
perl simuG.pl \
    -refseq SGDref.R64-2-1.fa.gz \
    -translocation_count 5 \
  -centromere_gff SGDref.R64-2-1.centromere.gff3 \
  -translocation_breakpoint_gff SGDref.R64-2-1.Ty1_FullLength.gff3 \
    -excluded_chr_list excluded_chr_list.yeast.txt \ # chrMT excluded
    -prefix yeast_test.TRA_run2 \
-seed 201812201903
```

Simulated human genome A (10000 SNPs):

```
perl simuG.pl \
    -refseq GRCh38.lite.fa.gz \
    -snp_count 10000 \
    -titv_ratio 2.0 \
    -excluded_chr_list excluded_chr_list.human.txt \ # chrM excluded
    -prefix human_test.SNP_INDEL \
-seed 201812201903
```

Simulated human genome B (1000 INDELs):

```
perl simuG.pl \
    -refseq GRCh38.lite.fa.gz \
    -indel_count 1000 \
    -excluded_chr_list excluded_chr_list.human.txt \ # chrM excluded
    -prefix human_test.SNP_INDEL \
    -seed 201812201903
```

Simulated human genome C (10 random CNVs due to segmental deletions):

```
perl simuG.pl \
    -refseq GRCh38.lite.fa.gz \
-cnv_count 10 \
-cnv_gain_loss_ratio 0 \
    -centromere_gff GRCh38.centromere.gff3 \
    -excluded_chr_list excluded_chr_list.human.txt \ # chrM excluded
    -prefix human_test.CNV_DEL \
    -seed 201812201903
```

Simulated human genome D (10 random CNVs due to dispersed duplications):

```
perl simuG.pl \
    -refseq GRCh38.lite.fa.gz \
-cnv_count 10 \
-cnv_gain_loss_ratio Inf \
-cnv_max_copy_number 3 \
-duplication_tandem_dispersed_ratio 0 \
    -centromere_gff GRCh38.centromere.gff3 \
    -excluded_chr_list excluded_chr_list.human.txt \ # chrM excluded
    -prefix human_test.CNV_DispersedDup \
    -seed 201812201903
```

Simulated human genome E (10 random CNVs due to tandem duplications):

```
perl simuG.pl \
    -refseq GRCh38.lite.fa.gz \
-cnv_count 10 \
-cnv_gain_loss_ratio Inf \
-cnv_max_copy_number 3 \
-duplication_tandem_dispersed_ratio Inf \
    -centromere_gff GRCh38.centromere.gff3 \
    -excluded_chr_list excluded_chr_list.human.txt \ # chrM excluded
    -prefix human_test.CNV_DispersedDup \
    -seed 201812201903
```

Simulated human genome F (5 random inversions):

```
perl simuG.pl \
    -refseq GRCh38.lite.fa.gz \
-inversion_count 5 \
-inversion_max_size 1000000 \
    -centromere_gff GRCh38.centromere.gff3 \
    -excluded_chr_list excluded_chr_list.human.txt \
    -prefix human_test.INV_run1 \
    -seed 201812201903
```

Simulated human genome G (5 random inversions with breakpoints sampled from full-length intact L1 transposable elements annotated in the reference genome):

```
perl simuG.pl \
    -refseq GRCh38.lite.fa.gz \
-inversion_count 5 \
-inversion_max_size 1000000 \
-centromere_gff GRCh38.centromere.gff3 \
-inversion_breakpoint_gff GRCh38.L1_FullLength.gff3 \
    -excluded_chr_list excluded_chr_list.human.txt \ # chrM excluded
    -prefix human_test.INV_run2 \
    -seed 201812201903
```

Simulated human genome H (5 random translocations):

```
perl simuG.pl \
    -refseq GRCh38.lite.fa.gz \
    -translocation_count 5 \
    -centromere_gff GRCh38.centromere.gff3 \
    -excluded_chr_list excluded_chr_list.human.txt \ # chrM excluded
    -prefix human_test.TRA_run1 \
    -seed 201812201903
```

Simulated human genome I (5 random translocations with breakpoints sampled from full-length intact L1 transposable elements annotated in the reference genome):

```
perl simuG.pl \
    -refseq GRCh38.lite.fa.gz \
    -translocation_count 5 \
  -centromere_gff GRCh38.centromere.gff3 \
  -translocation_breakpoint_gff GRCh38.L1_FullLength.gff3 \
    -excluded_chr_list excluded_chr_list.human.txt \ # chrM excluded
    -prefix human_test.TRA_run2 \
-seed 201812201903
```

The computational time used for each simulation is listed in Table S1, which was measured on a Linux computing server with an Intel Xeon CPU E5-2630 v3 (1.80 GHz) using a single thread.

| **Variant type** | **Variant number** | **Running time of simuG** | |
| --- | --- | --- | --- |
|  |  | **yeast genome** | **human genome** |
| SNP | 10000 | 3 s | 40 s |
| INDEL | 1000 | 1 s | 60 s |
| CNV: segmental deletion | 10 | 1 s | 40 s |
| CNV: dispersed duplication | 10 | 1 s | 40 s |
| CNV: tandem duplication | 10 | 1 s | 40 s |
| Inversion | 5 | 1 s | 40 s |
| Inversion with TE breakpoints | 5 | 1 s | 55 s |
| Translocation | 5 | 1 s | 50 s |
| Translocation with TE breakpoints | 5 | 1 s | 55 s |

Table S1. The design and computational time consumption of genome simulation with simuG. s: second.

3. **Illumina reads simulation**

For each simulated genome, we simulated 50X Illumina pair-end reads using ART v.MountRainier-2016-06-05 (Huang *et al.*, 2012) with the following parameters:

$art_dir/art_illumina \

--qprof1 $art_dir/Illumina_profiles/HiSeq2500L150R1.txt \ #profile

--qprof2 $art_dir/Illumina_profiles/HiSeq2500L150R1.txt \ #profile

-f 50 \ # fold of read coverage

-i <simulated_genome.fa> \

-l 100 \ # read length

-p \ # simulate paired reads

-na \ # no alignment output

-rs 201812 \

-m 500 \ # mean size of the paired-end fragment

-s 10\ # standard deviation of the fragment size

-o <output_prefix>

4. **PacBio reads simulation**

For simulated genome with CNV, inversions, and translocations, we simulated 25X PacBio reads using SimLoRd (Stöcker *et al.*, 2016) with the following parameters:

```
simlord_dir/simlord \
    --read-reference <simulated_genome.fa> \
    --coverage 25 \
    --no-sam \
    $prefix.simlord
```

5. **Illumina read mapping**

The simulated Illumina reads were trimmed by trimmomatic v0.38 (Bolger *et al.*, 2014) and then mapped to the corresponding reference genome (yeast or human) using BWA v0.7.17 (Li and Durbin, 2009). Samtools v1.9 (Li, 2011) and Picard tools v2.18.20 (<https://github.com/broadinstitute/picard>) were used for further processing (i.e. indexing, filtering, sorting, and duplicates removing) the resulting BAM files. Reads with mapping quality < 30 were discarded.

6. **PacBio read mapping**

The simulated PacBio reads were mapped to the corresponding reference genome (yeast or human) using minimap2 v2.16 (Li, 2018). Samtools v1.9 (Li, 2011) and Picard tools v2.18.20 (<https://github.com/broadinstitute/picard>) were used for further processing (i.e. indexing, filtering, and sorting) the resulting BAM files. Reads with mapping quality < 30 were discarded.

7. **SNP and INDEL variant calling and benchmarking**

For SNP and INDEL calling, we evaluated the performance of two widely used small variant callers: freebayes v1.2.0 (Garrison and Marth, 2012) and the HaplotypeCaller from GATK4 v4.0.11 (McKenna *et al.*, 2010) by applying them to the BAM file based on simulated reads from simulated genome A. We ran freebayes and GATK4’s HaplotypeCaller with default parameters with the only exception of setting the ploidy status to 1 (“-p” for freebayes and “-ploidy” for GATK4). The resulting VCF files from freebayes and GATK4 as well as the VCF file generated by simuG when simulating the SNP- and INDEL-bearing genomes were processed by vt (GitHub commit version vf6d2b5d) (Tan *et al.*, 2015) for variant decomposition, normalization, and annotation. The normalization step is very important for comparing VCF files generated from different methods since different tools might denote the same variant slightly differently in their respective VCF outputs depending on the immediate neighboring nucleotide bases of the corresponding variants. For each vt-processed VCF file, the SNP and INDEL variants were separated based on vt’s annotation and a final quality score cutoff of 30 was applied to filter out those low quality variants called by freebayes and GATK4. This step was performed by vcflib v1.0.0-rc2 (<https://github.com/vcflib/vcflib>). The true positive, false positive, and false negative value were further calculated by comparing the filtered VCF files from freebayes and GATK4 to the normalized simuG’s VCF output. Accordingly, precision, recall, and F_1_ score were further calculated using the following formula:

Precision = true positive/(true positive + false positive)

Recall = true positive/(true positive + false negative)

F_1_ score = 2 * (recall * precision )/(recall + precision)

8. **CNV, Inversion, and translocation calling and benchmarking**

For inversion and translocation calling, we evaluated the performance of the Illumina-read-based structural variant callers Delly v0.7.9 (Rausch *et al.*, 2012) and Manta v1.5.0 (Chen *et al.*, 2016) as well as the long-read-based structural variant caller Sniffles v1.011 (Sedlazeck *et al.*, 2018). We ran these tools with their default settings on the BAM files derived based on simulated genomes bearing CNVs, inversions, and translocations. The current version of Delly does not generate VCF file any more. So bcftools v1.9-83 (Li, 2011) was used to convert the BCF output from Delly to the VCF format. In contrast, Manta and Sniffles natively generate the VCF output. It is worth noting that different variant callers denote structural variants with different flavors of VCF formats. We manually examined and compared the VCF outputs from Delly, Manta, Sniffles, and simuG, during which we discarded those variants labeled as unresolved or low quality. Diagnostic measurements such as true positive, false positive, false negative, precision, recall, and F_1_ score were further calculated accordingly.
